# ACTIVE SUPER-ENHANCERS RELATED GENES PROFILING FOR THE IDENTIFICATION OF KEY ACTORS OF OSTEOSARCOMA METASTATIC DISSEMINATION

**DOI:** 10.64898/2026.11.25.769090

**Authors:** Mélanie Lavaud, Robel A. Tesfaye, Céline Charrier, Séverine Battaglia, Rose Anne Thépault, Marc Baud’huin, Bénédicte Brounais-Le Royer, Fernando Lecanda, Franck Verrecchia, François Lamoureux, Steven Georges, Benjamin Ory

## Abstract

We performed chromatin-immunoprecipitation coupled with sequencing (ChIP-Seq) against H3K27ac, a hallmark of active enhancers, on different osteosarcoma samples. Analysis of these generated datasets highlights distinct profiles of active super-enhancers (SEs), separating the osteosarcoma samples into two groups. This clustering of samples coincides with their metastatic capacities. Through RNA-Sequencing (RNA-Seq) data sets, we identified overexpressed target genes of active SEs, enclosed in their domains of action (Topologically Associated Domains), in metastatic samples. With the support of the literature, we highlight three genes that may play a crucial role in osteosarcoma metastatic evolution. Individual down expressions of these genes do not impair metastatic capacities of osteosarcoma cell lines in vitro, as opposed to the inhibition of the three identified genes simultaneously. Thus, through the study of the epigenome of various osteosarcoma samples, we highlighted that active SEs related genes can dictate tumour cells behaviour like their metastatic capacities. This is consistent with the observations of our peers demonstrating a crucial function of SEs in influencing cells fate, whether physiological or pathological. With this strategy, we dust off new robust therapeutic targets to fight against osteosarcoma, the most frequent malignant primary bone tumour.

## INTRODUCTION

Precise spatiotemporal regulation of gene expression is essential to determine cells fate and function. Using recent whole-genome analysis technologies, a variety of distant cis-acting DNA elements involved in the regulation of gene transcription have been identified and widely studied ^1^. Enhancers, first identified in the simian virus 40 genome ^2,3^, are periodic transcriptional thrusters and their activities are cell type specific ^4–8^. Clusters of enhancers, called super-enhancers (SEs), are more densely occupied by transcriptional activators than enhancers, resulting in higher expression of their distantly located target genes that have a prominent role in establishing and maintaining cellular identities ^9–11^. The activity of enhancers and SEs is restricted within isolated DNA domains, called Topologically Associated Domains (TADs), which exhibit strong and very frequent intra-domain contacts. The isolation of these TADs is achieved by the mechanism of chromatin loop extrusion, involving cohesin and CTCF molecules ^12,13^. Within each TAD, sub-chromatin loop structures allow physical proximity between enhancers/SEs and the promoters of their target genes, inducing transcriptional bursts ^14,15^. Since the discovery of SEs in 2013, different studies have highlighted the interest of genomic mapping of active SEs in order to identify genes involved in the establishment and maintenance of cell identity as well as in their behaviours, whether physiological or pathological.

Osteosarcoma is the most reported malignant bone tumour in children and young adults. At the time of diagnosis, 20% of patients present metastases, mainly in the lungs. Five years later, the survival rate of these patients is about 20%. Despite the overall improvement in the diagnosis and treatment of patients, the prognosis of some patients suffering from osteosarcoma remains unsatisfactory, especially if they have lung metastases ^16,17^. To develop new effective therapeutic strategies that prevent metastatic evolution of osteosarcoma, we need to better understand this malignant process. Our better understanding will allow us to uncover new therapeutic targets that have remained hidden until now. In the present study, we have tracked acetylations of H3K27 (lysine n°27 of H3 histones), a well-known epigenetic marker of active enhancers/SEs ^18^, by chromatin-immunoprecipitation coupled with sequencing (ChIP-Seq), in the genomes of osteosarcoma cell lines and primary osteosarcoma cells from diagnostic biopsies and surgical resections of lung metastases. In parallel, we conducted RNA-sequencing (RNA-Seq) to appreciate transcriptional programs of the mentioned osteosarcoma cells. A combined analysis of epigenetic and transcriptional profiles and an approach that considered the main characteristics of enhancers/SEs, allowed us to identify key genes involved in the metastatic evolution of osteosarcoma. Therefore, our approach consisting in focusing on active SEs allows us to identify novel markers involved in osteosarcoma metastatic evolution, which may be promising therapeutic targets to reduce the mortality of patients with osteosarcoma.

## MATERIAL AND METHODS

### Chromatin immunoprecipitation coupled with next-generation sequencing

Cells were harvested and washed with PBS (Eurobio, les Ulis, France). In order to cross-link proteins to DNA, washed cells were resuspended in 11% methanol-free formaldehyde (Thermo Fisher Scientific, Waltham, United States of America). Cross-linking was stopped by the addition of 125 mM glycine (Merck, Burlington, United States of America) and cells were washed twice with 1X EDTA-free protease inhibitor cocktail (PIC) (Sigma Aldrich, Saint Louis, United States of America). Then, cells in 1X EDTA-free PIC were snap frozen in liquid azote and sent to the Genomics Platform of the Broad Institute (Cambridge, United States of America). Chromatin immunoprecipitations (ChIP) were achieved using H3K27ac antibody (Cell Signaling Technology (8173S), Danvers, United States of America). Library preparation for 96 samples, comprising the ChIP samples used in this study, was performed in a multiplex manner on a 96-well plate. The multiplex strategy was chosen to remove the batch effect. Input sample library preparation and sequencing was performed at Active Motif Inc. (Carlsbad, United States of America) using a custom Illumina library type on an automated system (Apollo 342, Wafergen Biosystems/Takara). Prepared libraries for ChIP were sequenced using Illumina’s HiSeq 2500 in 2 x 100 b.p. format (San Diego, United States of America), and the input sample was sequenced using Illumina NextSeq 550 as 75 nucleotides long single-end reads.

### RNA next-generation sequencing

Cells were harvested and washed with cooled PBS. Dried-up cell pellets were snap frozen in dry ice and sent to Active Motif Inc. RNA sequencing (RNA-Seq) service. Libraries for RNA-SeqFsuper were prepared from purified RNA using Illumina TruSeq Stranded mRNA Sample Preparation kit. After library preparations, sequencing was done using Illumina’s NextSeq 500 in 2 x 42 bp format.

### Next-generation sequencing data treatments

#### ChIP-Sequencing data treatment

After aligning reads to the human genome (version hg19) using BWA (version 0.7.17), low-quality reads (-q 20) and PCR duplicates were discarded using samtools (version 1.9). To perform peak calling with MACS2, bedgraphs were built separately for the ChIP and Input samples based on the recommendations of MACS advanced manual. The Q-value of 10e-5 was set as the threshold for significant H3K27ac peaks. This threshold was used for all the samples and replicates in this study. The ROSE2 algorithm was applied on these significant peaks to identify SEs with default settings: peaks within 12500 bases windows were stitched together and those within 2500 bases of known TSS were excluded.

Identified SEs and significant peaks from all the osteosarcoma samples performed in duplicates were imported into R for further analyses. Reads overlapping the collected SEs were counted in each ChIP sample using RSubread’s featureCounts function. In order to take into account reads falling within overlapping SEs the argument allowMultiOverlap was set to true. This way reads within overlapping regions are counted once for each region rather than discarded due to ambiguity to assign them to a specific SE, as the function would perform by default. Then the over-estimation of reads in overlapping regions is corrected by keeping only the region with the highest read count using chromVAR’s filterPeaks function (v. 1.16.0). This means we consider the overlapping regions to be potentially the same SE. Biases in read counts of regions with equivalent GC% were estimated across samples using EDASeq (v. 2.28). Normalisation factors were then calculated to correct these biases and for each sample, the median of normalisation factors was extracted. Normalisation factors per sample were then exported and used to generate coverage files in bigwig format for consistent visualisation purposes with Deeptools bamCoverage (version 2.0). Sample normalisation factors from the object were used as scaleFactor to normalise coverage. Differentially activated regions between conditions were identified using Wald test (|log2 FC| > 1; BH adjusted p. value < 0.05).

#### RNA-Sequencing data treatment

Fastp was used to remove low-quality reads (-q 30) and remaining adapters. Filtered reads were then aligned to the human genome (hg19) using HISAT2 (version 2.1.0). Stringtie (version 2.1.1) was used to perform gene-level read quantification with a gtf from gencode (v. 35 hg19 liftover). Normalisation and differential expression analyses were performed using DESeq2 (v. 1.34.0). ClusterProfiler was used to perform Over Representation Analysis (ORA) and Gene Set Enrichment Analysis (GSEA) on selected gene sets (v. 4.2.2). Significantly differentially expressed genes were selected using Wald-test pairwise comparison (|log2 FC| > 1; BH adjusted p. value < 0.05).

#### Enhancer-promoter pairing

Enhancers/SEs target genes were identified using either the package inTAD (v. 1.14.0) or an in-house script. Correlation tests were performed using normalised H3K27ac signals of each regulatory element and the expression levels of genes found within the Topologically Associated Domain (TAD) of the regulatory element considered. TADs coordinates for human mesenchymal stem cells downloaded from the 3D browser were used in this effect. Correlations were considered significant when Pearson’s or Spearman’s tests output p.values below 0.05 and R2 or rho > 0.5. In order to capture enhancer-promoter pairs that have a functional effect, we filtered for pairs with genes that are significantly differentially expressed between conditions together with significantly differentially activated enhancers/SEs.

### Cells culture

Human HOS-MNNG, MG63, G292 and U2OS osteosarcoma cell lines were obtained from ATCC (Manassas, United States of America) and osteosarcoma primary cells were obtained from diagnostic biopsies (B: 531B and 652B) or from surgical resections of lung metastasis (M: 531MI and 531MII), performed on patients diagnosed with osteosarcoma at the Hospital of the University of Navarra (Clínica Universidad de Navarra, Pamplona, Spain). Samples were obtained following patients’ informed consent and after ethical approval by the Navarra University Hospital Ethics Committee. Cells, except 652B, were cultured in complete Dulbecco’s modified Eagle’s medium (DMEM, Eurobio) supplemented with 10% fetal bovine serum (FBS, Eurobio) at 37°C in humidified incubators containing 5% CO_2_ atmosphere. 652B were cultured in Minimum Essential Medium (MEM) Alpha (1X) (Thermo Fisher Scientific) supplemented with 10% FBS at 37°C in humidified incubators containing 5% CO_2_ atmosphere. In some experiences, osteosarcoma cells were treated with human recombinant IL31 (Thermo Fisher Scientific).

### Quantitative reverse transcription polymerase chain reaction (RT-qPCR)

Total RNA were extracted using NucleoSpin RNA Plus Mini Kit (Macherey-Nagel, Düren, Germany) and RNA concentrations were dosed using NanoDrop 1000 (Thermo Fisher Scientific). Reverse transcriptions were performed using Maxima H Minus First Strand cDNA Synthesis Kit (Thermo Fisher Scientific). Real-time monitoring of polymerase chain reactions was performed using preliminarily validated forward and reverse specific DNA primers (produced on request by Eurofins, Luxembourg, Luxembourg) and SYBR Select Master Mix (Thermo Fisher Scientific) in CFX96 Dx thermocycler (Bio-Rad, Hercules, United States of America). Forward and reverse specific DNA primers are provided in Table 1. Target genes expressions were normalised to B2M and HPRT genes expression levels and the relative expressions of target genes were calculated using the method described by Vandesompele et *al.*^19,20^.

**Table 1.**
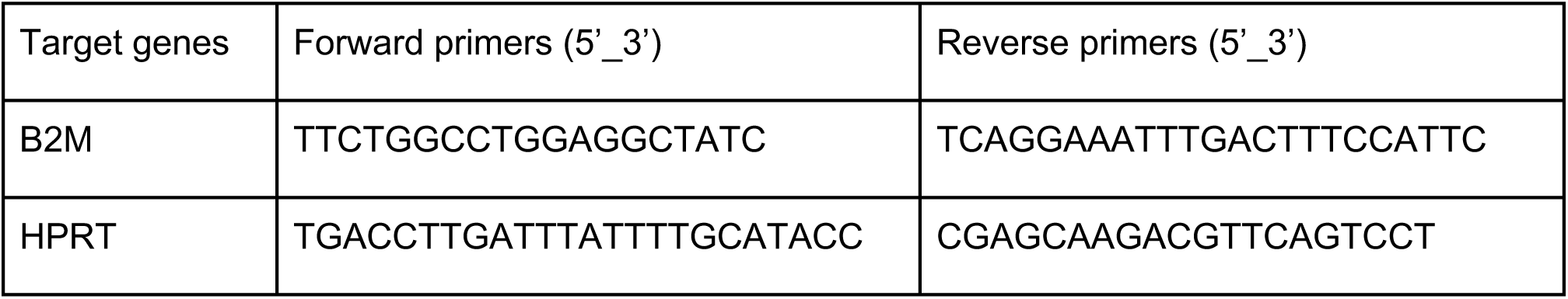
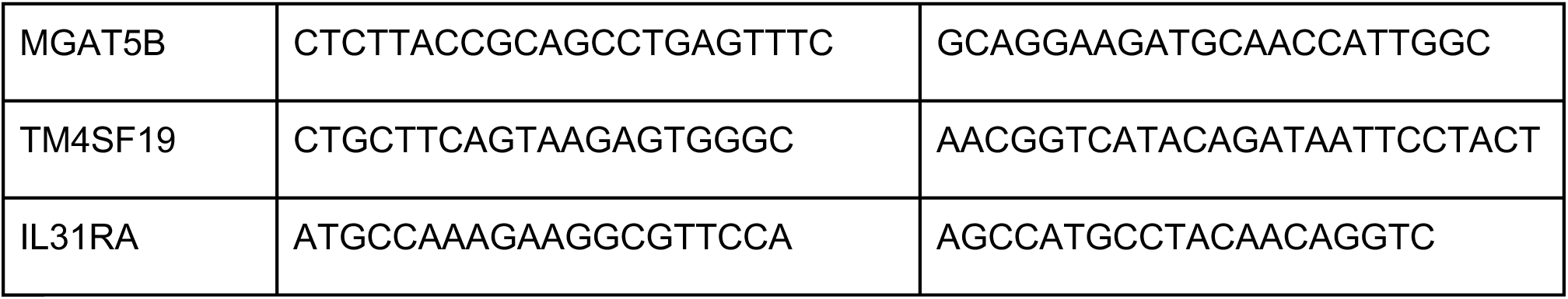
qPCR forward and reverse specific DNA primers.

### Western Blot

Total proteins were extracted using RIPA-based buffer containing protease and phosphatase inhibitors (Merck). Protein concentrations were dosed using BCA assay (Merck) and OD at 570 nm was measured with Victor 2 1420 multilabel counter (Perkin Elmer, Waltham, United States of America). Same amounts of whole protein extracts were prepared and reduced in 1X Bolt LDS Sample Buffer and 1X Bolt Reducing Agent (Thermo Fisher Scientific). Proteins were subjected to electrophoresis using precast polyacrylamide gels (Thermo Fisher Scientific) and transferred to methanol-activated polyvinylidene fluoride membranes (Merck). Membranes were blocked in Intercept Blocking Buffer (LI-COR, Lincoln, United States of America) and incubated with primary antibodies diluted in Intercept Blocking Buffer overnight at 4°C. Membranes were then washed intensively with TBS 0,1% Tween-20 (Merck) and incubated with secondary antibody, diluted in Intercept Blocking Buffer, for 1h at room temperature. Membranes were rewashed intensively with TBS 0,1% Tween-20 and proteins of interest were revealed using the Odyssey XF imaging system (LI-COR). Used antibodies (Cell Signaling Technology): 9145 (pSTAT3), 9139 (STAT3), 4650 (vinculin).

### Infiltration (transwell) assays

Cells were resuspended in DMEM supplemented with 1 % FBS and seeded on the upper side of inserts with porous transparent polyethylene terephthalate membranes (8µm pore size, Corning, Corning, United States of America). The lower chambers of the inserts were filled with DMEM 10 % FBS. Such 1 % - 10 % FBS gradient was generated between the upper and the lower sides of the inserts, to promote cells infiltration. Samples from cell resuspensions were seeded in 96-well plates in order to normalise the results of infiltration assays.

After 7 hours, cells on the upper side of the inserts were mechanically removed by using cotton swabs and cells that infiltrated the lower side were fixed to the membranes with 1 % glutaraldehyde (Sigma Aldrich) for 10 min at room temperature. After rinsing with PBS, cells were stained with 0.1 % crystal violet (Sigma Aldrich). Pictures of the infiltrated cells were taken using bright-field microscopy and they were counted using Fiji image processing package of ImageJ2 software (National Institutes of Health, Bethesda, United States of America). Cells seeded in 96-well plates were also fixed with 1 % glutaraldehyde, stained with 0.1 % crystal violet and, after drying, stainings were solubilized in Sorenson buffer (VWR, Radnor, United States of America). OD at 570 nm was measured with Victor 2 1420 multilabel counter (Perkin Elmer). Counts were normalised to corresponding OD values.

### Wound healing assays

Cell mats, seeded in 12-well plates in DMEM supplemented with 10% FBS, were wound using 1000µl pipette tips and rinsed with PBS before medium renewal and addition of mitomycin C (2.5µg/ml, Merck). Healings of cell mats were monitored using bright-field microscopy. Healings of cell mats were analysed using Fiji image processing package of ImageJ2 software.

### Individual genes knock downs

Individual knockdowns of target genes were realised by siRNA transfections, using Lipofectamine RNAiMAX reagent (ThermoFisher Scientific), purchased from Santa Cruz Biotechnology (Dallas, United States of America). siRNA against *GFP (Green fluorescent protein)* mRNA was used as control siRNA. RNAiMAX/siRNA used ratio was 1/2. siRNA final concentration was 30nM per well. siRNA transfections were performed 48h before presented assays.

### Simultaneous genes knock downs

Simultaneous knockdowns of target genes were realised by lentiviral transduction of Tet-On 3G tetracycline-inducible short hairpin RNAs (shRNAs) purchased from PerkinElmer (SMARTvector Inducible Lentiviral shRNA). Non-targeting shRNA was used as a control. A multiplicity of infection of 1 was used to transduce cells. In order to select transduced cells, cells were treated with 2,5 µg/ml puromycin (Sigma Aldrich) for 72 h and sorted by fluorescence-activated cell sorting assays (based on the expression of GFP induced by 1 µg/ml doxycycline (Sigma Aldrich) treatments for 72h). Expressions of shRNAs were induced by treatments of transduced cells with 1 µg/ml doxycycline for at least 72 h.

### Animal model

All experiments were conducted in Nantes medical school animal facility with the approval of the French Ministry of Agriculture for experimental use (Unité de Thérapeutique Expérimentale, Nantes; No. D44015). The protocol described in this project was approved by the local ethics committee CEE-PdL06 in Nantes (project APAFIS #8405). A paratibial injection with periosteal activation of 2×10^6^ HOS-MNNG cells was performed on 16 NMRI Nude mice (supplied by the company) and an injection into the caudal vein was performed on 8 NMRI Nude mice. The mice were anesthetized by inhalation of isoflurane (1.5%, 1L/min). Each mouse received modified HOS-MNNG cells (shRNAs MGAT5B 195, TM4SF19 636, IL31RA 878). Within each group, some of the mice received doxycycline (2 mg/mL) in their drinking water to enable transcription of the shRNAs. Doxycycline was dissolved in 5% water/sucrose. The water treatment was repeated three times a week. Weight, water intake, and tumor volume ((length x width x height)/2) were measured twice a week. The mice were sacrificed after 35 days of experimentation when the tumor volume reached 2000 mm3. The hind limbs and lungs were removed.

#### MicroCT analysis

The hind limbs of the mice were scanned using a high-resolution microCT scanner (Skyscan 1276, Bruker, Kontich, Belgium) with the following scanning parameters: source voltage 50kV, source current 200µA, 0.5 mm aluminum filter. Images were acquired with a pixel size of 17µm. Back projections were reconstructed using NRecon (Skyscan), and CTAn (Skyscan) software was used for segmentation and 3D analysis.

### Statistical analysis

Each experiment was conducted at least three times, independently, except for ChIP-Seq that have been performed once on two different samples. RNA-Seq assays have been performed once on three different samples. Cells transductions, selections and sorting were performed once. The results are presented as the means ± standard deviation. Mann–Whitney U test was used to compare two conditions in the GraphPad Prism 8 software (Dotmatics, Boston, United States of America). The differences were considered statistically significant at p. value < 0.05.

## RESULTS

### Active super-enhancers profiles segregate primary from metastatic osteosarcoma cells

Distant cis-acting DNA regulatory elements were identified in multiple osteosarcoma samples, including commercial cell lines (U2OS, HOS-MNNG, MG63, G292), primary tumour samples and metastatic samples derived from 2 patients (patients 652 and 531) by identifying significantly H3K27ac enriched regions. While we were only able to culture *in vitro* the primary sample from patient 652 (designated 652B), we were able to expand one primary sample and two samples from two metastases from patient 531 (designated as 531B, 531MI and 531MII respectively). ChIP-Seq against H3K27ac was performed in duplicates for each of these samples in a multiplexed manner to avoid the batch effect. A collection of active SEs in the different samples was obtained and read quantification was performed to evaluate the activation level of each regulatory element across the different samples. Whenever overlapping regions were encountered, only the highest scoring region was kept, making a total of 33921 active peaks, among which 3316 SEs. Efficient normalisation of read counts in the different samples was obtained by correcting biases in regions with equivalent GC% (**Figure supp 1 A**). After normalisation, hierarchical clustering and principal component analysis (PCA) on the SEs landscapes across the different samples display a separation between two sets of osteosarcoma samples (**Figure 1 A and 1 B, Figure supp 1 B and supp 1 C**). On the first principal component (PC1: 37% of variation explained), we observe a separation between the patient-derived metastatic samples and the patient-derived primary tumour samples. Along with these patient-derived tumour samples we observe that U2OS and HOS-MNNG seem to share closer active SEs landscapes to the patient-derived metastatic samples, while MG63 and G292 cluster with the primary samples. We also observe relatively similar active SEs landscapes between the primary samples from the 2 patients. On the other hand, the second PC (19% of variation explained) seemed to separate commercial cell lines from patient-derived samples except for U2OS. Interestingly, similar sample clustering was observed when PCA was performed on the transcriptomes with the 500 most variable genes (**Figure supp 1 D**).

**Figure 1.**
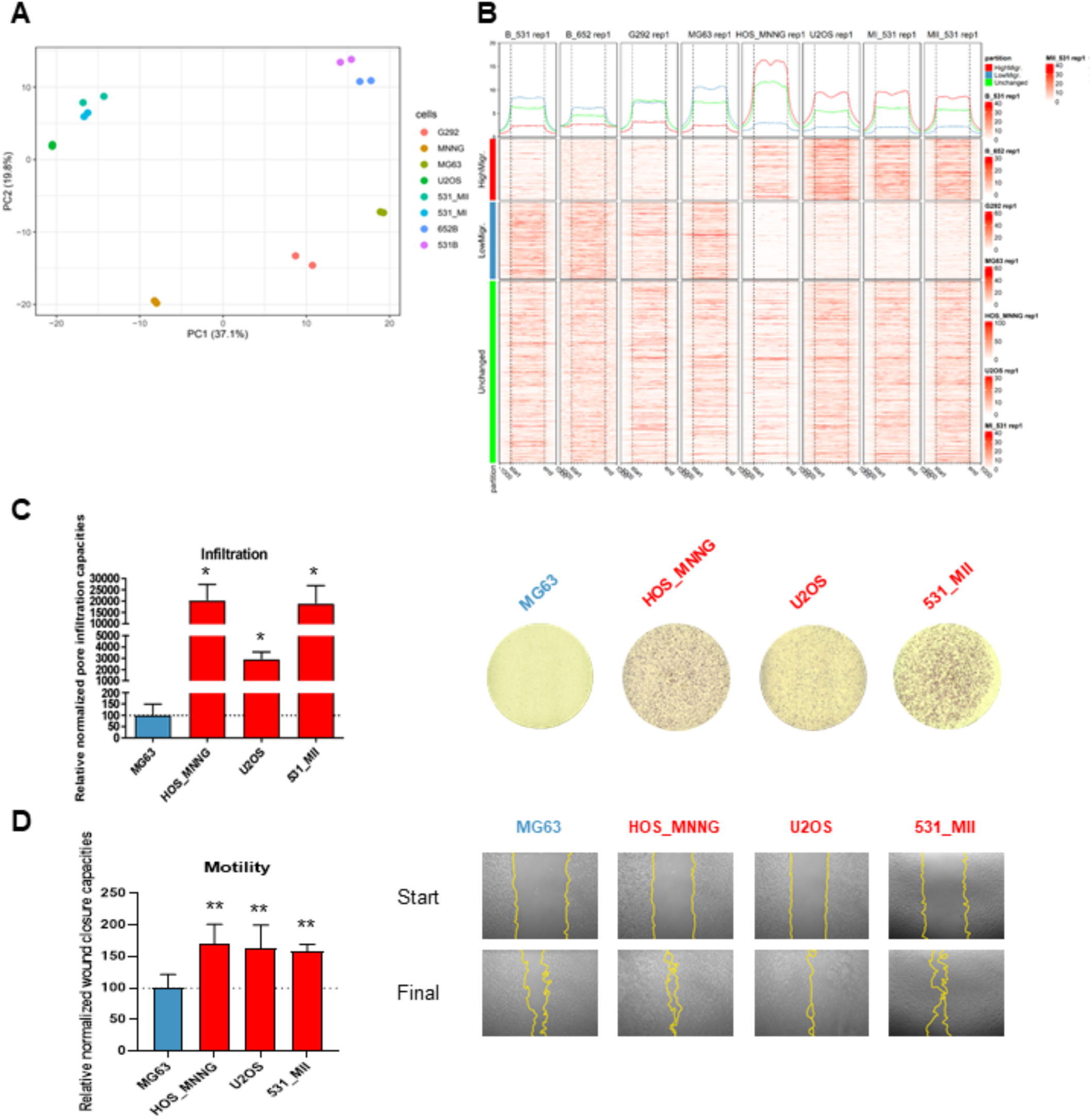
Active super-enhancers profiling predicts osteosarcoma metastatic capacity. (**A**) Principal Component Analysis (PCA) on all the super-enhancers identified in the different osteosarcoma samples considered in this study. (**B**) H3K27ac heatmap and average profiles within the groups of enhancers specific to each group of osteosarcoma samples identified from A. (**C**, **D**) Infiltration and motility capacities of osteosarcoma cells evaluated by transwell and wound healing assays respectively. Mann Whitney U test (one-tailed) p. value <0,05.

In order to validate the clustering of osteosarcoma samples, we evaluated the infiltration and motility capacities of the 2 groups of osteosarcoma samples (MG63 for the primary group; HOS-MNNG / U2OS / 531MII for the metastatic group) (**Figure 1 C and 1 D**). We observed better infiltration and motility capacities of osteosarcoma samples belonging to the metastatic group compared to the primary osteosarcoma cell line MG63. Infiltration capacities of cells clustered in the metastatic group are at least 30-fold more efficient than the primary sample. Motility capacities of HOS-MNNG, U2OS and 531MII osteosarcoma cell lines are approximately 1.5-fold higher than MG63. These results are consistent with the *in silico* observations based on the comparison of the epigenetic and transcriptional profiles of the osteosarcoma samples included in this study (**Figure 1 and Figure supp 1B,1D**).

### Active super-enhancers target genes identification highlights potential actors involved in metastatic evolution of osteosarcoma

Differentially enriched SEs were extracted between the two groups of osteosarcoma samples through a pair-wise comparison (Wald test; |log2 FC| > 1 and BH adjusted p. value < 0.05). In order to identify the target genes of differentially enriched SEs between the 2 groups of samples, correlation analyses were performed between the normalised H3K27ac enrichment in the SEs and the normalised expression levels of surrounding genes across the different samples (**Figure 2 A**). Genes enclosed within the TADs, as defined in mesenchymal stem cells, of the SEs of interest were considered for these correlation tests ^21,22^. Significant positive Spearman’s correlation (p. value < 0.05 and rho > 0.25) between a SE and expression of a gene located in the same TAD are the two criteria in this approach. With this enhancer-promoter pairing approach, we were able to associate one enhancer to multiple genes and one gene to multiple enhancers, which is in accordance with enhancers’ properties (**Figures supp 2 A**) ^23–25^.

**Figure 2.**
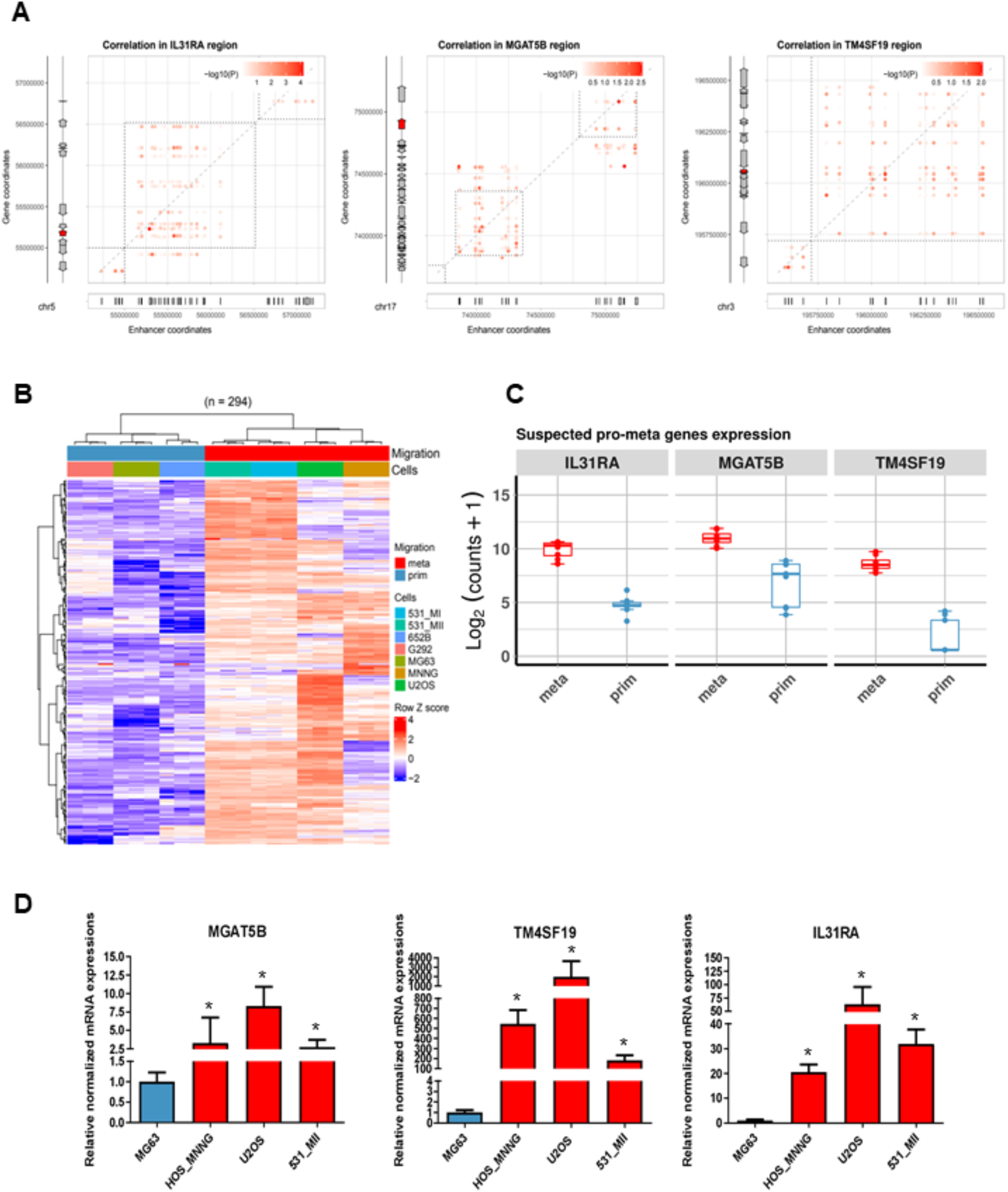
Prediction of target genes of metastatic group specific active super-enhancers. (**A**) Correlation analyses between enhancers normalized H3K27ac signals and expressions levels of genes found within the same TAD (Topologically Associated Domain). X-axis corresponds to genomic localizations of enhancers and Y-axis those of genes, TAD boundaries are presented in dashed grey lines. Colour code corresponds to significance of correlation (-log 10 (p. value)). (**B**) Heatmap of gene expression of predicted target genes of super-enhancers significantly more active in the metastatic group of osteosarcoma samples. (**C**, **D**) Expression levels of the 3 genes among the most up-regulated out of the potential targets of metastatic super-enhancers as measured by RNA-Sequencing and RT-qPCR respectively.

We identified 294 target genes of pro-metastatic active SEs that are also significantly up-regulated in their expression in the metastatic group (**Figure 2 B**). Hierarchical clustering on the expression levels of these target genes of metastatic SEs reveals a similar classification of osteosarcoma samples as the one obtained from the SEs profiles (**Figure supp 2 B**). Over-enrichment analysis on this list of genes indicates their involvement in osteoblasts development and in the regulation of the extracellular matrix (**Figures supp 2 C and supp 2 D**). After a literature study of the top ranking most up-regulated genes, we focused on 3 genes for posterior analyses: *MGAT5B*, *TM4SF19* and *IL31RA* (**Figure 2 C and supp 2 E**). Significant positive Spearman’s correlations were observed between these 3 genes expression levels and the associated SEs normalised H3K27ac enrichment across the considered osteosarcoma samples (**Figure supp 2 F**). Track plots displaying H3K27ac enrichment within the SEs targeting these genes show higher enrichment in the metastatic group than in the non-metastatic group of osteosarcoma samples (**Figure supp 2 G and supp 2 H**). GnT-Vb, encoded by *MGAT5B*, is an enzyme that controls the synthesis of complex cell surface glycans and has been reported to be overexpressed in human metastatic hepatocellular carcinoma (HCC) cell lines compared to human non-metastatic HCC cell line (Hep3B) ^26^. GnT-V and GnT-Vb products were revealed to be promising markers and targets of both metastatic competence in mice models and disease-free survival in patients in mammary cancer and prostate cancer respectively ^27,28^. Similarly, overexpression of *TM4SF19* increases metastatic colonisation in melanoma mice models ^29^. *IL31RA* overexpression was reported to promote tumour cell motility in breast cancer *in vitro* and silencing *IL31RA* expression decreases pulmonary metastasis development in animal models ^30^. RNA-Seq analyses and RT-qPCR on biological replicates indicate significant overexpression of these 3 genes in the metastatic group compared to the primary group (**Figures 2 C and 2 D**). We performed IL31 treatments of osteosarcoma samples to evaluate IL31RA responsiveness to these treatments by measuring STAT3 and pSTAT3 levels. As expected, the MG63 cell line does not respond to IL31 treatments compared to HOS-MNNG, U2OS and 531MII (**Figure supp 2 I**).

### Simultaneous down expressions of identified super-enhancers-related genes decrease the metastatic propensity of osteosarcoma cells *in vitro*

Our observations along with the literature allowed us to suspect the involvement of the 3 identified genes in the metastatic evolution of osteosarcoma. In order to validate their implication in this process, we performed artificial down expressions of each gene using siRNAs followed by functional assays. After validation of knockdowns of our identified genes (**Figure 3 A**), we performed transwell assays. Our results demonstrate that individual down expressions of the 3 identified genes do not significantly impair the infiltration capacity of HOS-MNNG cell line, despite an anti-infiltrative tendency of approximately 25-30% when *IL31RA* and *MGAT5B* are knocked down (**Figure 3 B**). Wound healing assays also demonstrate that knocking down the 3 potential pro-metastatic genes does not decrease the motility capacity of the HOS-MNNG cell line (**Figure 3 C**). To understand these results, we examined expressions of the 3 genes for each individual down expression. Surprisingly, we observed that knocking down *MGAT5B* expression seems to increase the expression of *TM4SF19* and vice versa. Furthermore, down expressions of *MGAT5B* and *TM4SF19* also may impair *IL31RA* expression. *IL31RA* expression knockdown seems to increase the expression of *TM4SF19* (**Figure supp 3 A**). Thus, inhibition of one target gene could result in the establishment of a compensation system in which the other two identified genes are overexpressed.

**Figure 3.**
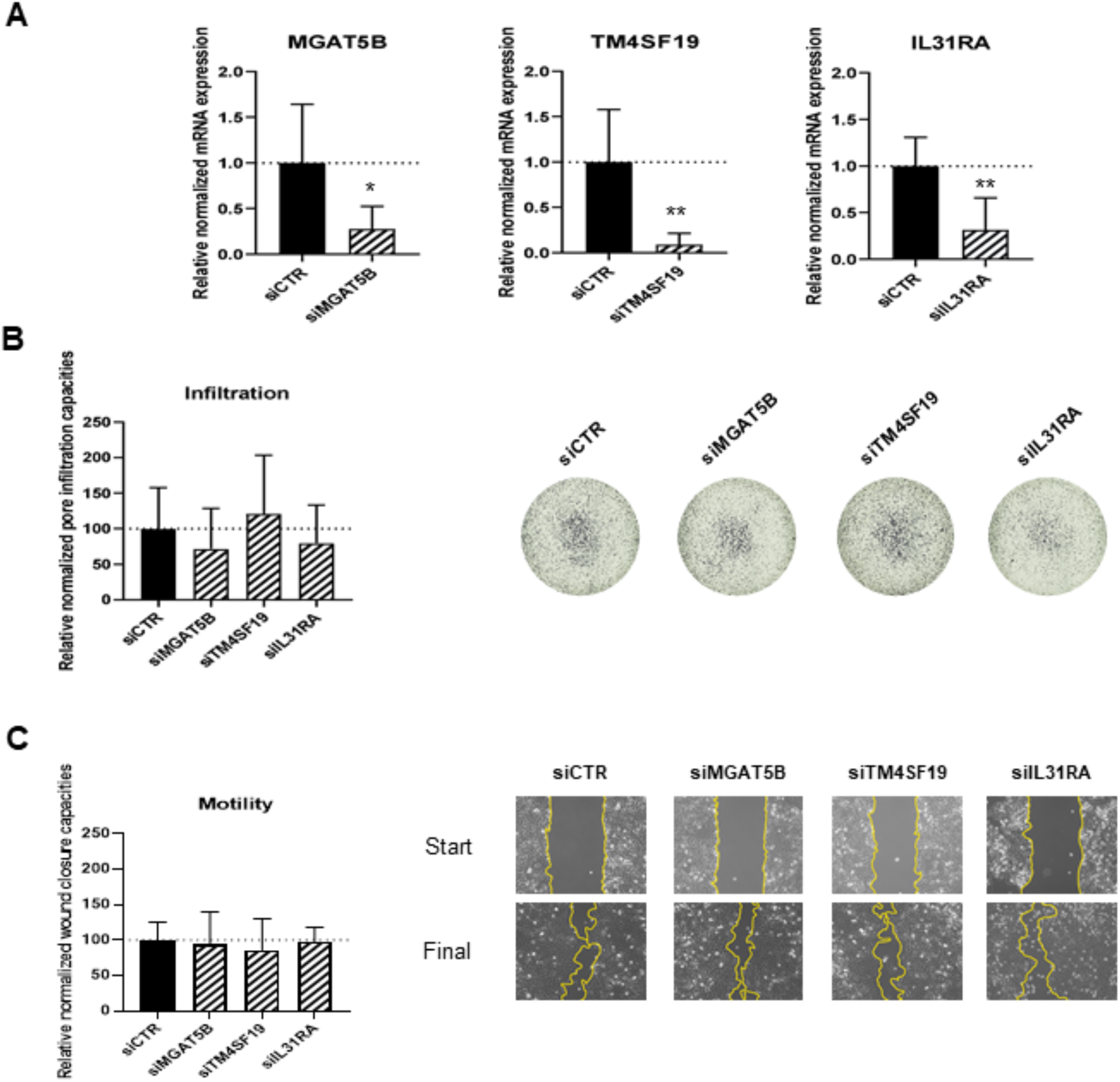
*In vitro* individual down expressions of identified active super-enhancers related genes do not impair infiltration and motility capacities of osteosarcoma cells. (A) Validations by RT-qPCR of individual target genes knock downs realized by HOS-MNNG siRNA transfections. (B, C) Infiltration and motility capacities of HOS-MNNG presenting individual target genes knock downs evaluated by transwell and wound healing assays respectively. Mann Whitney U test (one-tailed) p. value <0,05.

In order to cope with this potential compensatory effect, we aimed to knock down the expressions of the 3 identified genes simultaneously. We performed simultaneous knock downs of our 3 identified genes by performing shRNAs transduction (2 different sets of shRNAs, **Figure 4 A**, **4 D and Figure supp 4 A**). Simultaneous down expressions of the 3 metastatic SE-induced genes decrease infiltration and motility capacities of HOS-MNNG cell line as estimated by transwell and wound healing assays respectively (**Figure 4 B and 4 C**, **Figure 4 E and F**).

**Figure 4.**
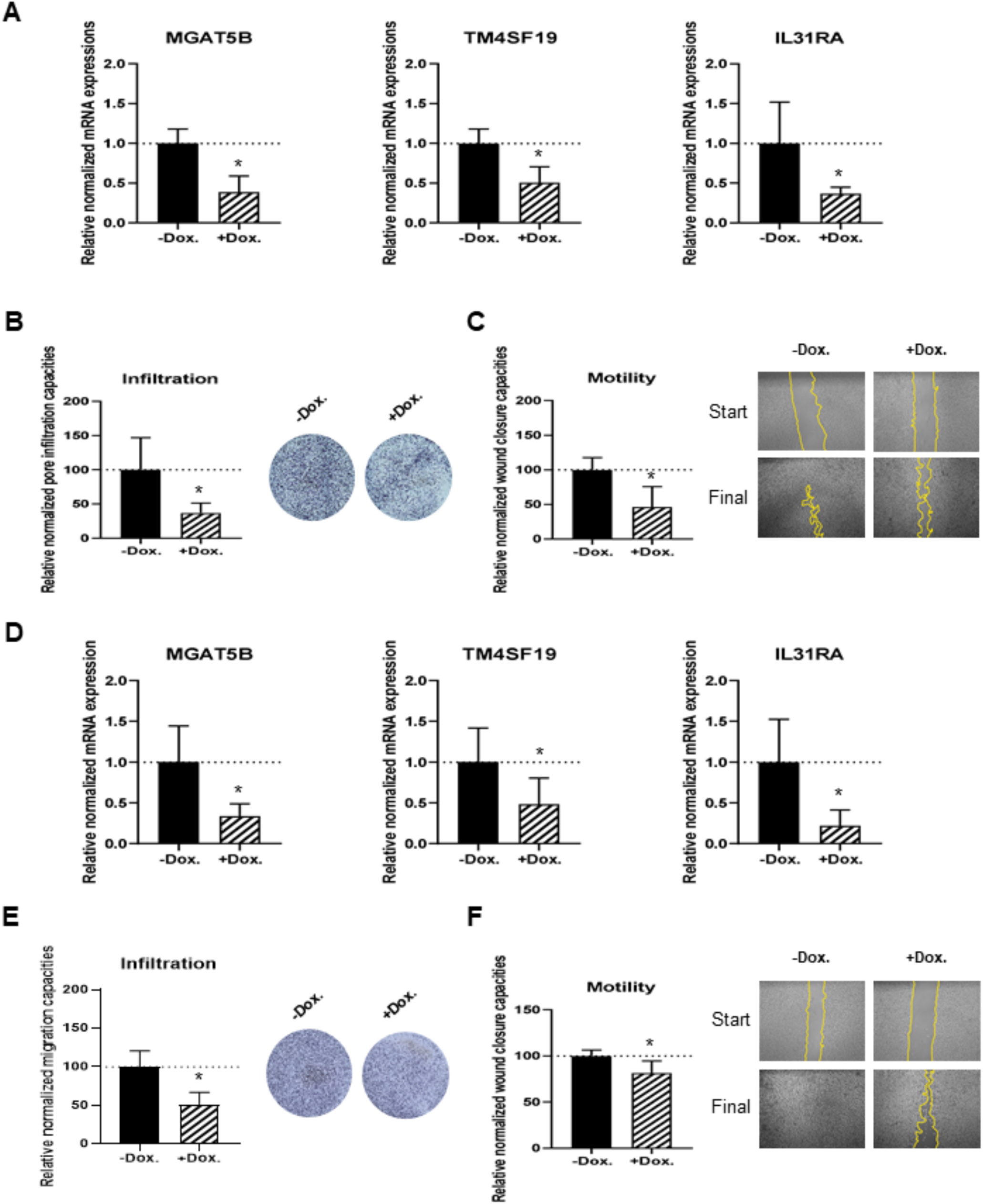
*In vitro* simultaneous down expressions of identified active super-enhancers related genes impair infiltration and motility capacities of osteosarcoma cells. (A) Validations by RT-qPCR of target genes simultaneous knock downs realized by HOS-MNNG triple shRNA transductions (shRNAs 195 636 878). (**B**, **C**) Infiltration and motility capacities of HOS-MNNG presenting simultaneous target genes knock down evaluated by transwell and wound healing assays respectively (shRNAs 195 636 878). (**D**) Validations by RT-qPCR of target genes simultaneous knock downs realized by HOS-MNNG triple shRNA transductions (shRNAs 794 012 446). (**E**, **F**) Infiltration and motility capacities of HOS-MNNG presenting simultaneous target genes knock down evaluated by transwell and wound healing assays respectively (shRNAs 794 012 446). Mann Whitney U test (one-tailed) p. value <0,05.

## DISCUSSION

Analysis of the epigenetic and transcriptomic profiles allowed efficient separation of the osteosarcoma samples included in this study on the basis of their *in vitro* motility (wound healing assays) and infiltration (transwell assays) capacities. Hierarchical clustering on the profiles of the most H3K27ac enriched epigenetic elements displays comparative epigenomes between patient-derived metastatic tumour extracts and well-known commercial osteosarcoma cell lines. Interestingly, metastatic and primary site tumour extracts from the same patient (patient 531) display distinct SEs profiles and cluster within separate groups. This indicates that the epigenetic and transcriptomic profiles activated in the metastatic group samples could be common across osteosarcoma samples. However, this observation requires further confirmation by including in the study other metastatic and primary patient-derived osteosarcoma samples. The complex genetics observed in osteosarcoma could cause diverse epigenetic and transcriptional mechanisms for metastatic evolution.

Functional assays on the infiltrative and motility capacities of the osteosarcoma samples also confirmed the *in silico* classification. We designated the group containing the tumour samples from metastases of osteosarcoma patients, as the metastatic group. Admittedly, *in vitro* assays are approximations to measure the metastatic potentials of mainly the commercial cell lines, and mice osteosarcoma models using these commercial cell lines should confirm this classification.

In order to associate more precisely SEs to their target genes, correlation tests between the regulatory regions and genes within their reach were performed. This allowed identifying genes that demonstrate transcriptional alterations along with the activation states of their potential regulatory DNA sequences. We focused on differentially activated SEs across the 2 groups of osteosarcoma samples to capture SE-promoter pairs with differential activities in the 2 groups. This choice of highlighting differentially activated SEs allowed to exclude correlations with insignificant effects on the transcriptomes of the osteosarcoma samples.

Validating functionally the *in silico* predictions of targets of the pro-metastasis SEs could be time-consuming, as this requires individual efficient silencing of the 294 target genes and evaluating the effect of the gene silencing on metastatic potential in mice models. We chose to work on the 3 genes presented in this article because they displayed consistently strong up-regulations when metastatic group individual samples were compared with the primary group individual samples. They were also supported by the literature mainly from studies on other cancers. A CRISPR-CAS9-based screening, targeting either the SEs or the target genes, followed by *in vitro* infiltration and motility capacities assessment could be a more feasible strategy to perform a more comprehensive analysis.

Furthermore, *in vivo* experiments are indispensable to fully validate the involvement of the 3 identified genes in the metastatic evolution of osteosarcoma, as this allows to recapitulate all the different steps enabling metastases development from primary tumour. Thus, osteosarcoma cells presenting simultaneous down expressions of *MGAT5B*, *TM4SF19* and *IL31RA* should be used to create an osteosarcoma mouse model and to evaluate lung metastases development.

---

**Figure supp 1.**
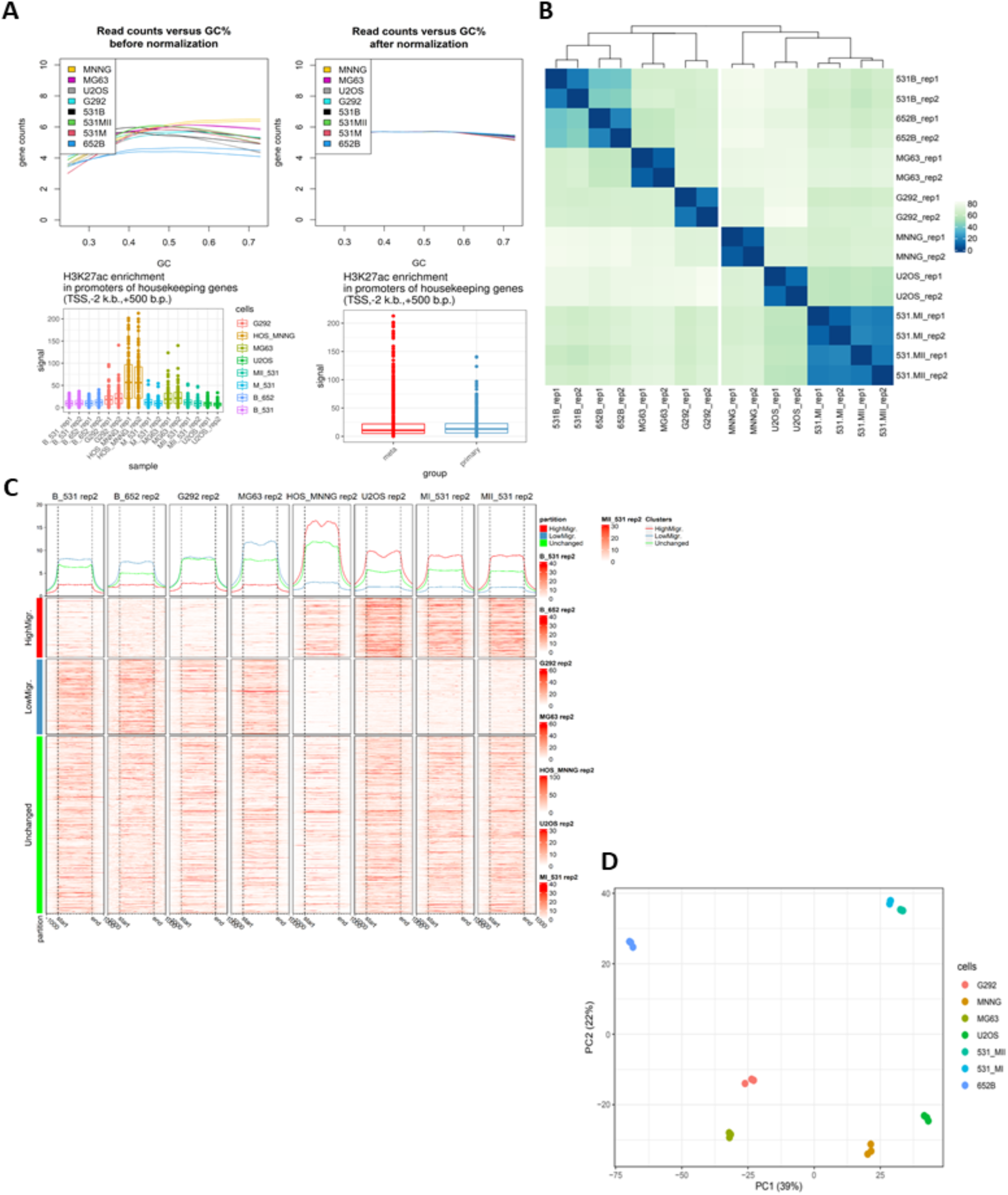
Active super-enhancers profiling predict osteosarcoma metastatic capacity. (**A**) Checking normalization process: Read counts across samples within regions of equivalent GC% before and after normalization with EDASeq (top) and H3K27ac enrichment after normalization in the promoters of 96 housekeeping genes per sample and per group (bottom). (**B**) Hierarchical clustering based on normalized read counts in the active super-enhancers of osteosarcoma samples. (**C**) H3K27ac heatmap and average profiles within the groups of enhancers specific to each group of osteosarcoma samples identified from Figure 1 A. (**D**) Principal Component Analysis (PCA) on the 500 most variable genes across the transcriptomes of the different osteosarcoma samples considered in this study.

**Figure supp 2.**
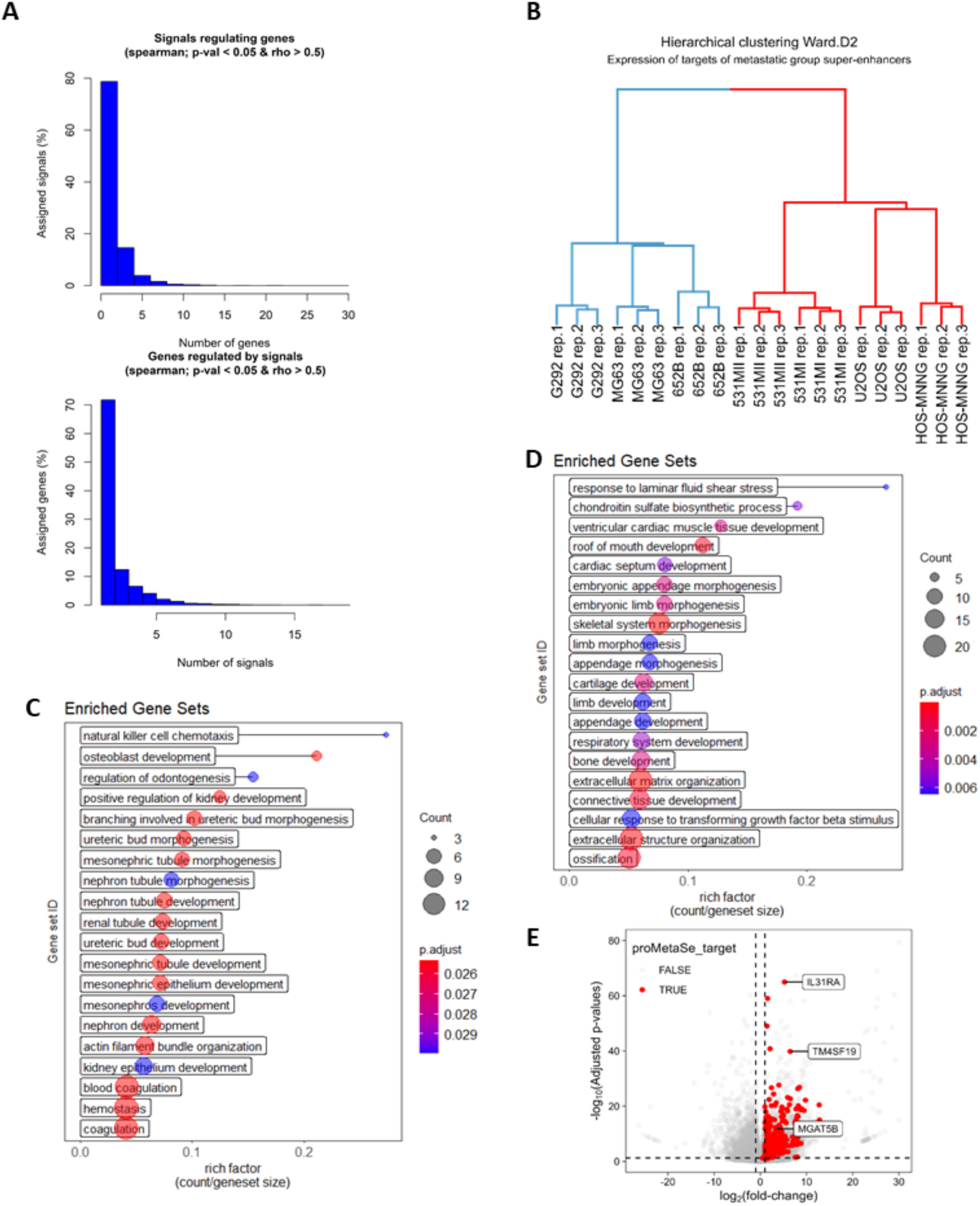
Prediction of target genes of metastatic group specific active super-enhancers. (**A**) Number of genes assigned per enhancer (top) and number of enhancers associated to a gene (bottom). (**B**) Hierarchical clustering based on the expression of targets of metastatic group specific super-enhancers. (**C**) Over-enrichment analysis (ORA) with GO:BP on the list of potential targets of metastatic super-enhancers (from Figure 2 B). (**D**) Over-enrichment analysis (ORA) with Reactome.db on the list of potential targets of metastatic super-enhancers (from Figure 2 B). (**E**) Volcano plot in the comparison transcriptomes of « metastatic » group samples and « primary » group samples. Predicted targets of metastatic group specific super-enhancers are plotted in red.

**Figure supp 2.**
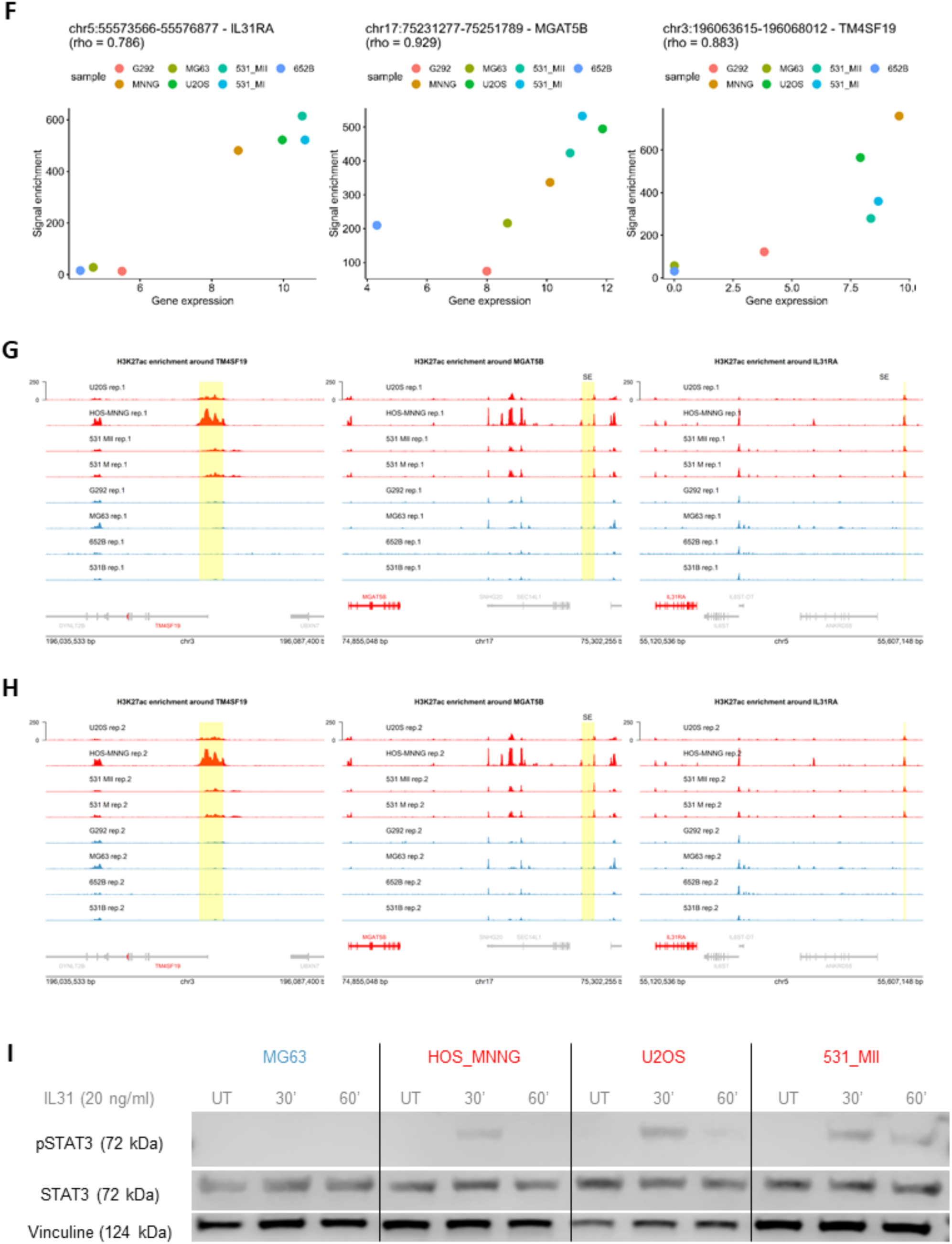
Prediction of target genes of metastatic group specific active super-enhancers. (**F**) Spearman correlation tests between super-enhancers normalised H3K27ac signal and predicted target gene expression across osteosarcoma samples. (**G**, **H**) H3K27ac trackplot around predicted target genes of super-enhancers (super-enhancers are highlighted in yellow) across first and second replicates of osteosarcoma samples respectively. (**I**) STAT3 and pSTAT3 proteins expression under IL31 treatments (20 ng/ml) performed during 30 or 60 minutes (UT: untreated).

**Figure supp. 3.**
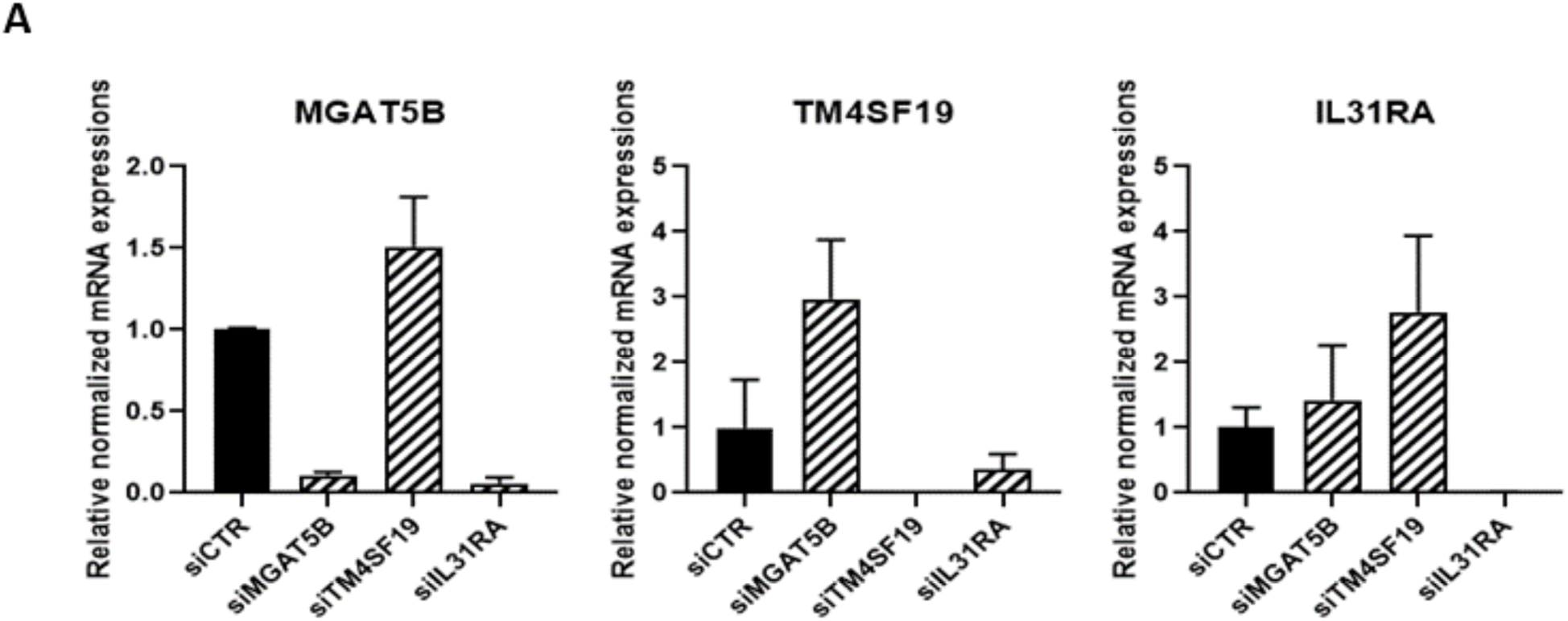
*In vitro* individual down expressions of identified active super-enhancers related genes do not impair infiltration and motility capacities of osteosarcoma cells. (A) Evaluation by RT-qPCR of identified genes expression under individual target genes knock downs realized by HOS-MNNG siRNA transfections.

**Figure supp. 4.**
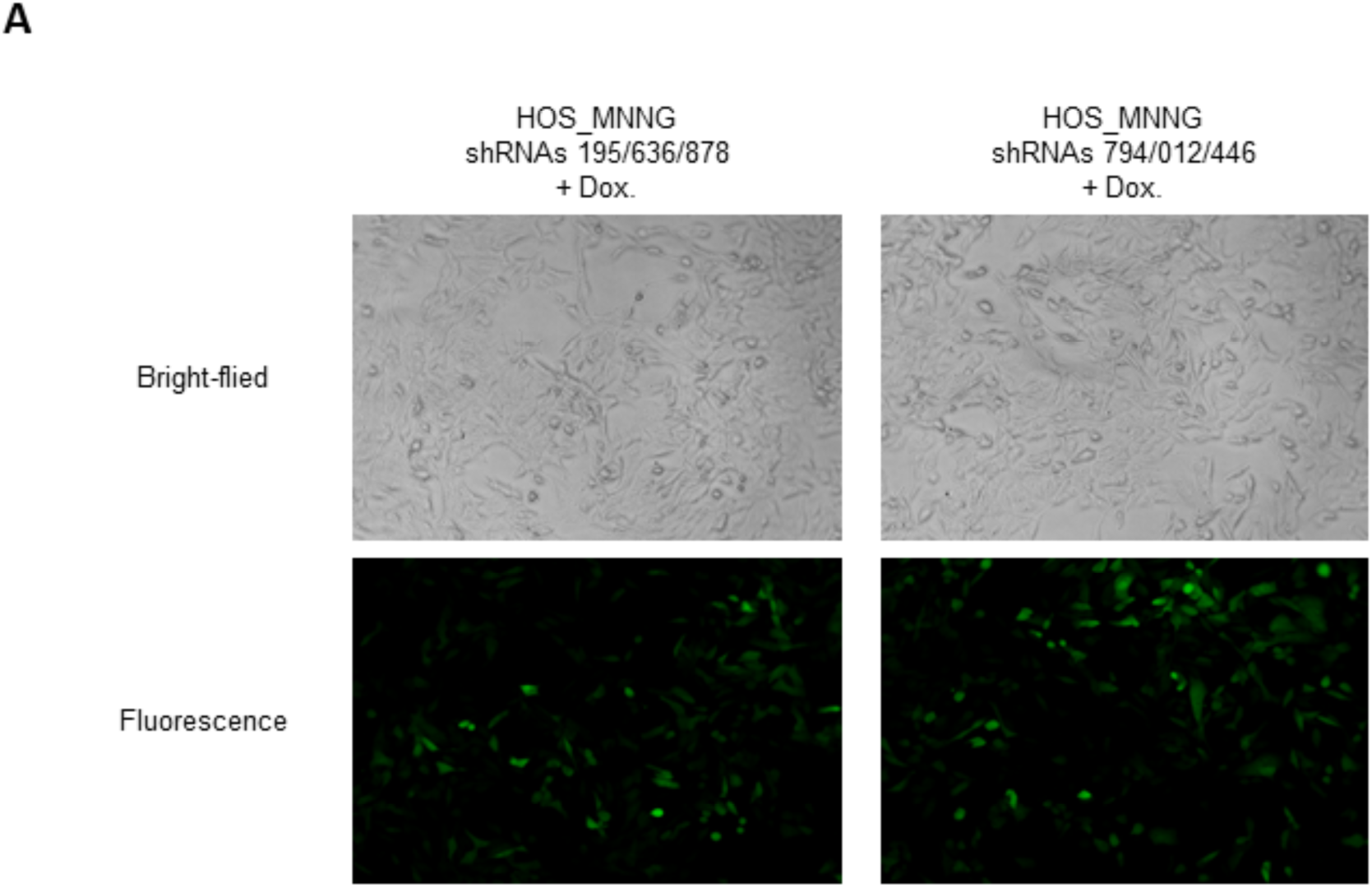
*In vitro* simultaneous down expressions of identified active super-enhancers related genes impair infiltration and motility capacities of osteosarcoma cells. (**A**) Induction of shRNAs expressions by 1µg/ml doxycycline treatments (72h) also activates Green Fluorescent protein expression.

